# AntiCP 2.0: An updated model for predicting anticancer peptides

**DOI:** 10.1101/2020.03.23.003780

**Authors:** Piyush Agrawal, Dhruv Bhagat, Manish Mahalwal, Neelam Sharma, Gajendra P. S. Raghava

**Author notes:** **Emails of Authors:** PA, DB, MM, NS, GPSR. **Corresponding author** Prof. G.P.S. Raghava, Head of Department, Department of Computational Biology, Indraprastha Institute of Information Technology, Okhla Phase 3, New Delhi-110020, India., E-mail address.

## Abstract

Increasing use of therapeutic peptides for treating cancer has received considerable attention of the scientific community in the recent years. The present study describes the *in silico* model developed for predicting and designing anticancer peptides (ACPs). ACPs residue composition analysis revealed the preference of A, F, K, L and W. Positional preference analysis revealed that residue A, F and K are preferred at N-terminus and residue L and K are preferred at C-terminus. Motif analysis revealed the presence of motifs like LAKLA, AKLAK, FAKL, LAKL in ACPs. Prediction models were developed using various input features and implementing different machine learning classifiers on two datasets main and alternate dataset. In the case of main dataset, ETree Classifier based model developed using dipeptide composition achieved maximum MCC of 0.51 and 0.83 AUROC on the training dataset. In the case of alternate dataset, ETree Classifier based model developed using amino acid composition performed best and achieved the highest MCC of 0.80 and AUROC of 0.97 on the training dataset. Models were trained and tested using five-fold cross validation technique and their performance was also evaluated on the validation dataset. Best models were implemented in the webserver AntiCP 2.0, freely available at https://webs.iiitd.edu.in/raghava/anticp2. The webserver is compatible with multiple screens such as iPhone, iPad, laptop, and android phones. The standalone version of the software is provided in the form of GitHub package as well as in docker technology.

## Introduction

Cancer is the second most dangerous disease, leading to deaths globally, after cardiovascular diseases. According to WHO report, in 2017, 9.56 million people died prematurely due to cancer, worldwide, which states that every 6^th^ person dying in the world is because of cancer. The detection of cancer at an early stage is one of the major challenges. If the cancer is diagnosed at an early stage, then there are higher possibilities of surviving and less morbidity. However, the lack of early diagnosis of cancer is one of the major barriers in treating the patients [1]. Due to the inadequacy of accurate and non-invasive markers, the detection of cancer is usually biased [2]. Recent advancements in the field of genomics and proteomics have led to the discovery of peptide-based biomarkers which has enhanced the detection of cancer at an early stage [3]. After diagnosing cancer, the next step involves its treatment. Currently, chemotherapy, radiation therapy, hormonal therapy, and surgery are the conventional treatments available for treating cancer. Adverse side effects and high cost of these conventional methods are the obstacles for effective treatment [4], and even if the treatment is successful, then there are the chances of reoccurrence of cancer [5], which indicates towards the need of better and more effective treatment. In the past few years, peptide-based therapy has emanated as an advanced and novel strategy for treating cancer [6]. It has several advantages like high target specificity, good efficacy, easily synthesized, low toxicity, easily modified chemically [7], less immunogenic when compared to recombinant antibodies [8]. In recent years, therapeutic peptides have appeared as a diagnostic tool and have the ability to treat many diseases [9–14]. More than 7000 naturally occurring peptides have been reported in the last decade, which exhibit multiple bioactivities (antifungal, antiviral, antibacterial, anticancer, tumor-homing, etc.) [15]. As per the report, more than 60 drugs have been approved by the FDA, and >500 are under clinical trials [16].

Anticancer peptides (ACPs) are part of the antimicrobial peptide group which exhibits anticancer activity. These are small peptides [5-50 amino acids] and cationic in nature. Mostly they possess α-helix as the secondary structure (e.g. LL-37, BMAP-27, BMAP-28, Cercopin A, etc.) or folds into β-sheet (e.g. Defensins, Lactoferrin, etc.). Some peptides have shown extended linear structure such as Tritrpticin and Indolicidin [17,18]. Cancer cell exhibit different properties in comparison to normal cells. Cancer cells possess larger surface area due to presence of a higher number of microvilli, negative charge of the cell membrane, higher membrane fluidity, etc. [19–21]. These features allows cationic anticancer peptides to interact with the negatively charged membrane via electrostatic interactions ultimately leading to necrosis i.e. selective killing of cancer cells [22]. Other means by which ACPs exhibits its function includes lysing of mitochondrial membrane (apoptosis), inhibiting angiogenesis pathway or recruiting other immune cells for attacking cancer cells, and activating essential proteins which ultimately lyse cancer cells [23]. Different mechanism of ACP function is shown in **Figure 1**. In order to explore the mechanism of action and development of novel therapeutic ACPs, accurate prediction of ACPs is very essential. As the experimental process is time consuming, labour intensive and costly, there is need for computational tools to do the same. In past, many sequence-based methods have been proposed for predicting and designing ACPs. Some of the popular methods include AntiCP [24], iACP [25], ACPP [26], iACP-GAEnsC [27], MLACP [28], SAP [29], TargetACP [30], ACPred [31], ACPred-FL [32], PTPD [33], Hajisharifi et. al’s method [34], Li and Wang’s method [35]. Detail information for most of these methods is provided in an article by Schaduangrat et al. [31].

**Figure 1:**
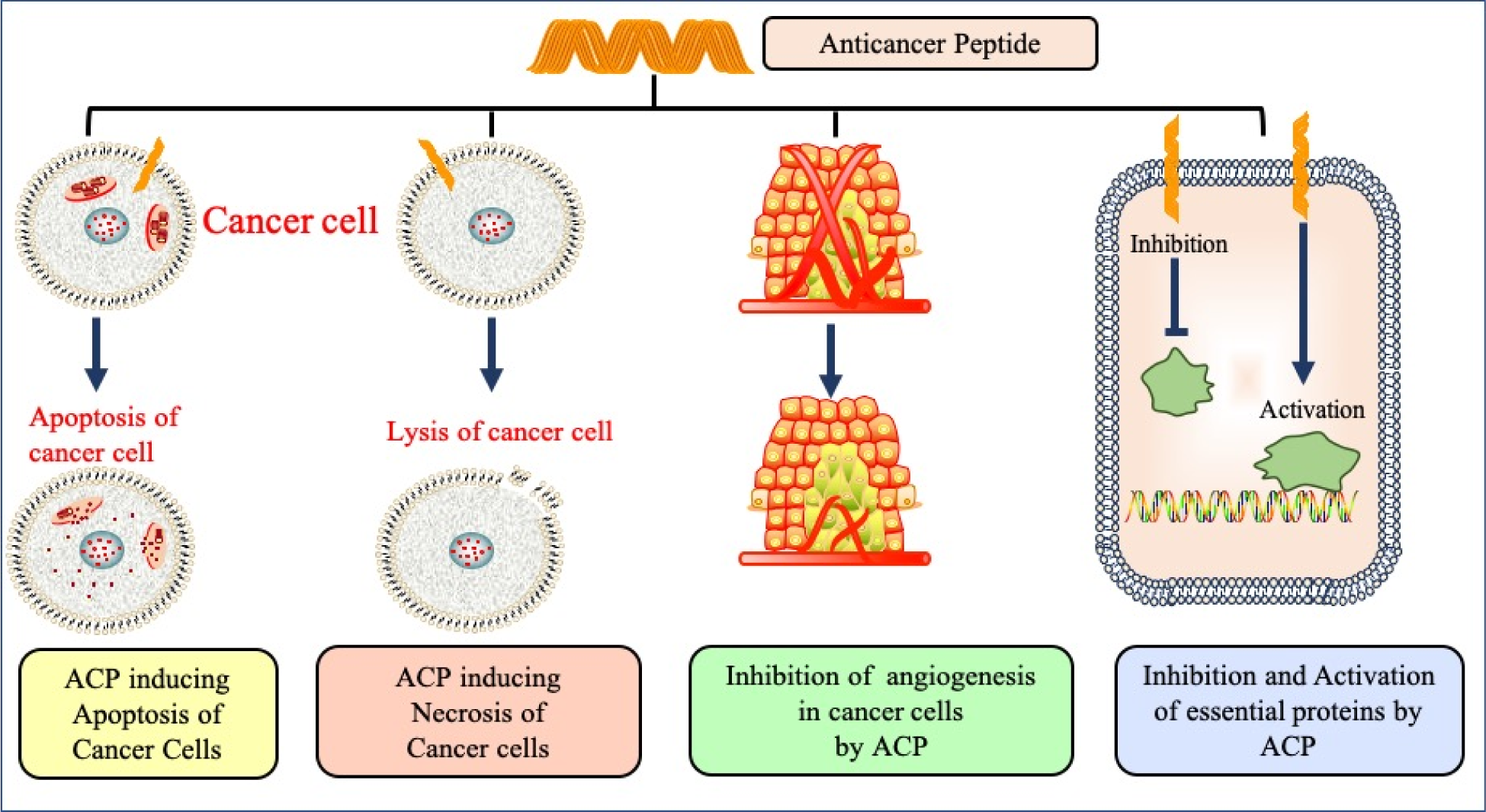
Various mechanism of action of anticancer peptides.

However, there are certain limitations associated with the methods mentioned above. The major limitation includes the selection of dataset both quality and quantity wise for developing the respective methods. This creates a challenging situation for an experimental scientist in selecting the method and the feature for the prediction study. In addition, some of the methods such as iACP-GAEnsC and TargetACP do not provide webserver facility, hence, limiting the utility of these methods for biologists. The other important drawback of the above mentioned methods is their inability to discriminate peptides having similar composition but different activity. Aforementioned issues motivated us to come up with a method which addresses the above mentioned limitations. In the current study, we have developed a method AntiCP 2.0. The method is developed on the biggest dataset reported till to date. The performance of the model was also evaluated on the datasets of the similar peptide with different activity i.e. ACPs and non-ACPs.

## MATERIALS AND METHODS

### Datasets Preparation

We created two balanced datasets, (i) main dataset and (ii) alternate dataset. The two datasets consisted of peptides, which consists of only natural amino acids and lengths varying from 1 to 50. Detail description of the dataset creation used in the study is given below:

i. **Main dataset:** In the main dataset, experimentally validated ACPs are taken as positive class and AMPs, which did not show any anticancer properties were considered as negative class. Initially, the dataset comprises of consists of 970 peptides; however, some of the peptides were found to be common in ACPs and non-ACPs. Therefore, to avoid biasness during model development, we removed such peptides and were finally left with 861 unique peptides.
ii. **Alternate dataset:** In the alternate dataset, the number of peptides in the positive and negative classes was 970, as there were no common peptides. Here, the experimentally validated ACPs were considered as a positive class, and for negative class, random peptides were generated using SwissProt proteins. This kind of approach has been used earlier [36,37].

### Internal and External Validation

The datasets were randomly divided into two parts (i) training dataset, which comprises of 80% data and (ii) validation dataset with remaining 20% data. In case of internal validation, we developed prediction models and evaluated them using five-fold cross validation technique. In five-fold cross validation, sequences are divided randomly into five datasets, out of which any four datasets are used for training and the fifth one is used for testing purposes. To use each of the five datasets at least once for testing, we repeat this process 5 times. The final result is calculated by averaging the performance of all five sets. For external validation, we evaluated the performance of the model developed using training dataset on validation dataset.

### Additional dataset

We created an additional dataset and compared the performance of developed models on it. The additional dataset comprises of the peptides which are very similar in amino acid composition. This exercise was performed because, in the past it has been shown that it is challenging to discriminate similar peptides with different activity [38]. For creating this dataset, the euclidean distance was computed in between the ACPs and non-ACPs, and peptides with minimum distance were selected. For this, we considered main dataset. This is a well-established approach in the literature and has been used previously [39,40].

### MERCI Motifs Analysis

MERCI (Motif-EmeRging and with Classes-Identification) software [41] was used for searching the motifs uniquely present in anticancer peptides. We used the default parameters for running the software. Motif analysis renders the information related to various kind of patterns, which could be present in the anticancer peptides. Default parameters were used for running the MERCI code.

### Input features for prediction

We used multiple input features and for developing prediction models, we applied various machine learning techniques. The various features used are described below:

#### (a) Amino acid composition-based model

Residue composition provides us insight about the fraction of amino acid type present within the peptide. AAC gives the information regarding the percentage of each residue present in the protein/peptide, which is a vector of dimension 20. The formula for calculating AAC for each amino acid residue is illustrated in equation. The formula for calculating AAC for each amino acid residue is illustrated in equation (1)

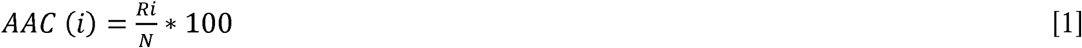

where, AAC (i) is the amino acid per cent composition (i); R_i_ is the number of residues of type i, and N represents the length of the peptide sequence.

#### (b) Dipeptide composition-based model

Dipeptide composition (DPC) was also used as an input feature. It provides not only the composition of the pair of the amino acid but also their local order with a vector of dimension 400 (20 × 20). To compute the DPC of the given protein/peptide, we first compute all the possible pairs of the 20 amino acids, also known as dipeptides (e.g. A-A, A-C, A-D…Y-W, Y-Y), and then compute their composition. The formula for calculating DPC is illustrated in equation (2)

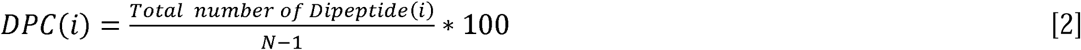

where, DPC (i) is a type of dipeptide out of 400 dipeptides and N is the length of the peptide.

#### (c) Split composition

We have also computed both the composition (amino acid and dipeptide) for 5, 10 and 15 residues present at N and C terminus of the protein/peptide. We have also combined the terminal residues like N5C5, N10C10 and N15C15 and then again computed the composition.

#### (iv) Binary profile-based model

Since the length of peptides used in this study is variable, it is challenging to generate a fixed-length pattern. To overcome this, we have extracted the segments of fixed-length from either N-terminus or C-terminus of the peptide to generate a fixed-length binary profile [40]. After computing the fixed-length patterns, we have generated the binary profiles for the residues of both the terminus, i.e., N5, N10, N15, C5, C10, C15 and similarly for combined terminal residues N5C5, N10C10 and N15C15.

### Machine learning techniques

Various machine learning techniques were implemented in this study using the scikit-learn package [42]. A brief description of the package is provided below.

Scikit-learn is a free software machine learning library for the Python programming language. It features various classification, regression and clustering algorithms including support vector machines (SVM). We used 6 machine learning classifiers from this package namely Support Vector Machine (SVM), Random Forest (RF), k-Nearest Neighbours (KNN), Extra Trees (ETree), Artificial Neural Network (ANN) and Ridge Classifier. We tuned different parameters present in these classifiers during the run and reported the results obtained on the best parameters.

### Performance measure

We measured the performance of our methods using threshold dependent and threshold independent parameters. Threshold dependent parameters includes Sensitivity (Sen), Specificity (Spc), Accuracy (Acc) and Matthews Correlation Coefficient (MCC). These parameters are calculated using the following equations:

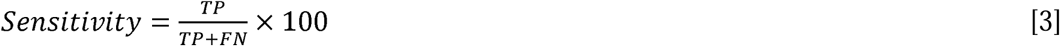

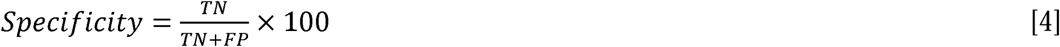

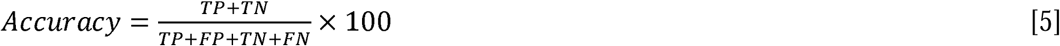

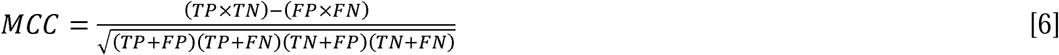

where TP represents the correct positive predictions, TN represents the correct negative predictions, FP represents the false positive predictions, which are actually negative, and FN represents the false negative predictions, which are actually positive. For the threshold independent parameter evaluation, Area Under Receiver Operating Characteristics (AUROC) curve was calculated where a ROC curve was drawn in between false positive and false negative rates.

## Result

Various types of analysis were performed such as composition analysis, residue preference analysis, and exclusive motifs present in the peptides. For the analysis, we used main dataset. Results of these analysis is explained below.

### Composition analysis of ACPs and non-ACPs

Percent average of amino acid composition was computed using the ACPs and non-ACPs of the main dataset. ACPs were found to be rich in residues like A, F, K, L, and W when compared to non-ACPs and proteins present in SwissProt **(Figure 2)**. Non-ACPs were found to be rich in residues like C, G, R, and S. Comparison with residue composition of SwissProt proteins was made to check that the residue composition of ACPs is not random and can be easily differentiated with normal proteins/peptides. The analysis shows that ACPs are rich in positively charged residue and aromatic amino acids. The positively charged residues of the ACPs interacts with the negatively charged cancer cell membrane and carry out its lysis.

**Figure 2:**
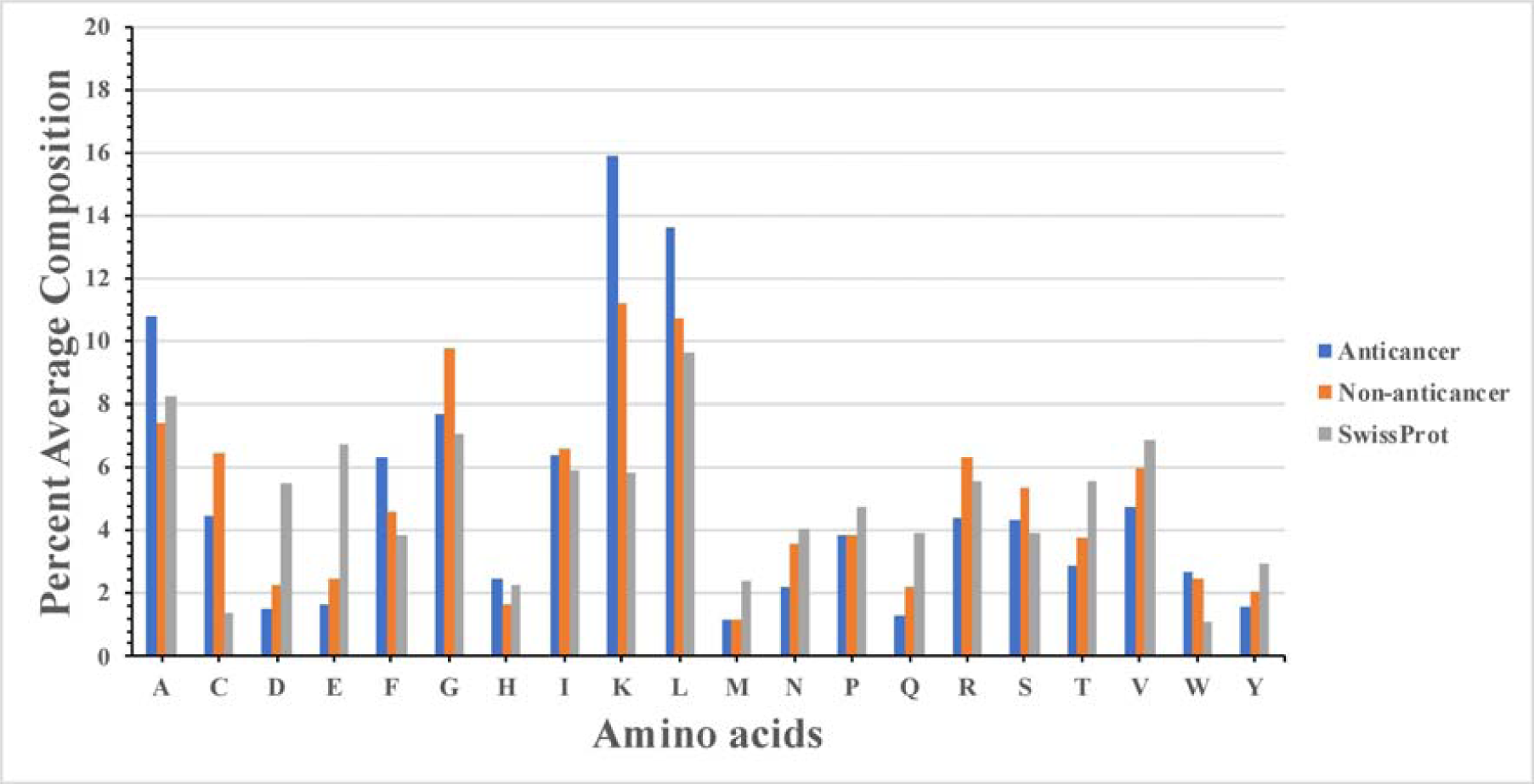
Percentage composition analysis of residue present in ACPs, non-ACPs and SwissProt proteins.

### Residue Preference

Two sample logo was computed to understand the residue preference at the termini (N and C). The first 10 residues were taken for generating the logo from both termini. As shown in **Figure 3A**, at N-terminus, preference of residue F was higher at 1^st^ position, A at 2^nd^ position, and K was dominating at 3^rd^ position. Apart from these residues, L was also preferred at other positions. At C-terminus, residue L and K were found to be highly preferred in comparison to other amino acids **(Figure 3B)**.

**Figure 3:**
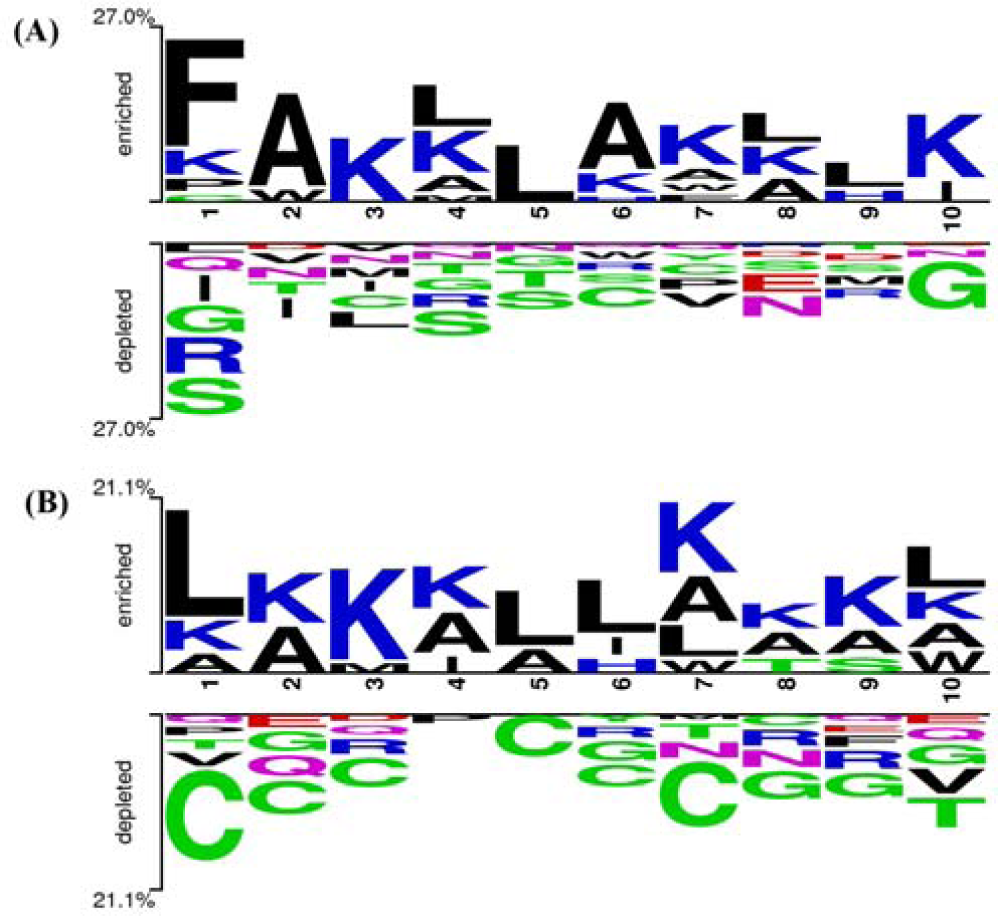
Two Sample Logo explaining the positional preference of first 10 residue present in ACPs and non-ACPs at (A) N-terminus and (B) C-terminus respectively.

### Motif Analysis

Exclusive motifs present in ACPs and non-ACPs were extracted using MERCI software. In the case of the main dataset, we found that motifs such as “LAKLA, AKLAK, FAKL, LAKL” were present in ACPs and motifs like “GLW, CKIK, DLV, AGKG” were present in non-ACPs. Exclusive motifs frequently found in ACPs and non-ACPs are provided in **Supplementary Table S1**.

### Machine learning model performance on various features

In the last few decades, in silico identification and designing of therapeutic peptides has been in trend. Several *in silico* based methods have been developed regarding the same, eventually helping biologists to screen potential molecules in lesser time and cost. These in silico models have been developed based on features extracted from the experimental data. These features are crucial for the proper functioning of the ACPs. Amino acid composition, dipeptide composition, terminus residue, motif information, binary profile, physiochemical properties are some of those essential features which differentiate one class of molecule with others. In this study too, prediction models have been developed using these features. The prediction models were developed by implementing classifiers present in the scikit-learn package. In the current study, we have implemented several classifiers (See Material & Methods section) for developing prediction models. The performance of these classifiers on various input features is provided below.

### Model Performance on Main Dataset

Residue composition has been a key feature in classifying one class of molecules with other. Hence prediction models were developed using this feature on both datasets. In the case of the main dataset, ETree based model performed best with 74.78% sen, 74.06% spc, 74.72% acc, 0.49 MCC, and 0.82 AUROC on the training dataset. On validation dataset, it achieved 79.19% sen, 68.79% spc, 73.99% acc, 0.48 MCC, and 0.83 AUROC **(Table 1)**. For dipeptide composition too, ETree based model performed best with 74.06% sen, 76.52% spc, 75.29% acc, 0.51 MCC, and 0.83 AUROC on the training dataset. On validation dataset, it achieved 77.46% sen, 73.41% spc, 75.43% acc, 0.51 MCC, and 0.83 AUROC **(Table 2)**.

**Table 1.** The performance of amino acid composition based models developed using different machine learning techniques on main dataset.

| Techniques<br>(Parameters) | Training dataset |  |  |  |  | Validation dataset |  |  |  |  |
| --- | --- | --- | --- | --- | --- | --- | --- | --- | --- | --- |
|  | Sen | Spc | Acc | MCC | AUROC | Sen | Spc | Acc | MCC | AUROC |
| SVC<br>(g=0.001, c=1) | 72.90 | 68.99 | 70.94 | 0.42 | 0.78 | 73.41 | 65.32 | 69.36 | 0.39 | 0.79 |
| RF<br>(Ntree =100 ) | 74.35 | 73.33 | 73.84 | 0.48 | 0.82 | 78.03 | 68.21 | 73.12 | 0.46 | 0.83 |
| ETree<br>(Ntree =400) | 74.78 | 74.06 | 74.42 | 0.49 | 0.82 | 79.19 | 68.79 | 73.99 | 0.48 | 0.83 |
| MLP<br>(activation=logistic,<br>solver=lbgfs,<br>hidden_layers=17,<br>max_iter=10) | 63.91 | 69.86 | 66.88 | 0.34 | 0.73 | 61.85 | 69.94 | 65.90 | 0.32 | 0.71 |
| KNN<br>(neighbors=10,<br>algorithm=brute,<br>weights=uniform) | 67.54 | 70.29 | 68.91 | 0.38 | 0.76 | 71.10 | 69.36 | 70.23 | 0.40 | 0.78 |
| Ridge (alpha = 0) | 67.83 | 57.97 | 62.90 | 0.26 | 0.70 | 69.36 | 58.96 | 64.16 | 0.28 | 0.71 |

**Sen:** Sensitivity, **Spc:** Specificity, **Acc:** Accuracy, **MCC:** Matthews Correlation Coefficient, **AUROC:** Area Under the Receiver Operating Characteristic curve

**Table 2.** The performance of dipeptide composition based models developed using different machine learning techniques on main dataset.

| Techniques<br>(Parameters) | Training dataset |  |  |  |  | Validation dataset |  |  |  |  |
| --- | --- | --- | --- | --- | --- | --- | --- | --- | --- | --- |
|  | Sen | Spc | Acc | MCC | AUROC | Sen | Spc | Acc | MCC | AUROC |
| SVC<br>(g=0.001, c=1) | 75.94 | 68.84 | 72.39 | 0.45 | 0.81 | 75.72 | 66.47 | 71.10 | 0.42 | 0.80 |
| RF<br>(Ntree =100 ) | 75.07 | 74.06 | 74.57 | 0.49 | 0.83 | 80.92 | 67.05 | 73.99 | 0.48 | 0.83 |
| ETree<br>(Ntree =400) | 74.06 | 76.52 | 75.29 | 0.51 | 0.83 | 77.46 | 73.41 | 75.43 | 0.51 | 0.83 |
| MLP<br>(activation=logistic,<br>solver=lbgfs,<br>hidden_layers=17,<br>max_iter=10) | 69.71 | 69.13 | 69.42 | 0.39 | 0.78 | 75.72 | 68.79 | 72.25 | 0.45 | 0.80 |
| KNN<br>(neighbors=10,<br>algorithm=brute,<br>weights=uniform) | 72.46 | 69.13 | 70.80 | 0.42 | 0.79 | 76.88 | 67.63 | 72.25 | 0.45 | 0.81 |
| Ridge (alpha = 0) | 70.43 | 69.42 | 69.93 | 0.40 | 0.75 | 71.68 | 70.52 | 71.10 | 0.42 | 0.76 |

SVC based models were also developed using the split amino acid composition for various patterns (N5, N10, N15, C5, C10, C15, N5C5, N10C10, and N15C15). The model developed using the N5 pattern performed best with 0.46 MCC and 0.81 AUROC on the training dataset and 0.42 MCC and 0.78 AUROC on the validation dataset. A detailed result for other patterns is provided in **Supplementary Table S2.** Likewise, in the case of SVC based models developed using split dipeptide composition, the model developed using the N5 pattern performed best. It achieved the highest MCC of 0.49 and AUROC of 0.86 on the training dataset and MCC of 0.51 and AUROC of 0.83 on the validation dataset. The result for other patterns is provided in **Supplementary Table S3.**

We also developed SVC models utilizing the binary profile of peptide. The binary profile is an important feature when it comes to classifying two classes of peptides, as shown in previous studies [43]. Models were developed for different lengths of peptides (first 5, 10, and 15) from N and C-terminus. Also, models were developed by joining the peptides obtained from both N and C terminus i.e., N5C5, N10C10, and N15C15. In the case of the main dataset, the model developed using the N10C10 pattern performed best among all models. On training dataset, sen of 76.32%, spc of 68.61%, acc of 72.44%, MCC of 0.45, and AUROC of 0.81 was obtained. On the validation dataset, the model achieved sen of 79.39%, spc of 66.27%, acc of 72.81%, MCC of 0.46, and AUROC of 0.81. The result for all other patterns for the main dataset is provided in **Table 3.**

**Table 3.** The performance of SVC based model developed on main dataset, where models were developed using binary profile of part of peptide.

| Techniques<br>(Parameters) | Training dataset |  |  |  |  | Validation dataset |  |  |  |  |
| --- | --- | --- | --- | --- | --- | --- | --- | --- | --- | --- |
|  | Sen | Spc | Acc | MCC | AUROC | Sen | Spc | Acc | MCC | AUROC |
| N5<br>(g=0.01, c=4) | 73.56 | 66.83 | 70.03 | 0.40 | 0.77 | 73.72 | 66.46 | 70.06 | 0.40 | 0.76 |
| N10<br>(g=0.01, c=4) | 73.57 | 66.56 | 70.04 | 0.40 | 0.79 | 77.50 | 64.42 | 70.90 | 0.42 | 0.80 |
| N15<br>(g=0.01, c=4) | 70.09 | 68.14 | 69.00 | 0.38 | 0.78 | 73.45 | 66.91 | 69.84 | 0.40 | 0.76 |
| C5<br>(g=0.01, c=4) | 64.10 | 63.25 | 63.66 | 0.27 | 0.68 | 61.82 | 66.87 | 64.33 | 0.29 | 0.70 |
| C10<br>(g=0.01, c=4) | 68.71 | 63.17 | 65.92 | 0.32 | 0.73 | 73.01 | 58.54 | 65.75 | 0.32 | 0.74 |
| C15<br>(g=0.01, c=4) | 66.52 | 61.69 | 63.83 | 0.28 | 0.72 | 67.83 | 63.12 | 65.23 | 0.31 | 0.74 |
| N5C5<br>(g=0.01, c=4) | 69.76 | 68.65 | 69.20 | 0.38 | 0.77 | 72.35 | 70.41 | 71.39 | 0.43 | 0.79 |
| N10C10<br>(g=0.01, c=4) | 76.32 | 68.61 | 72.44 | 0.45 | 0.81 | 79.39 | 66.27 | 72.81 | 0.46 | 0.81 |
| N15C15<br>(g=0.01, c=4) | 73.76 | 70.98 | 72.22 | 0.44 | 0.80 | 72.17 | 65.96 | 68.75 | 0.38 | 0.79 |

\* **Sen:** Sensitivity, **Spc:** Specificity, **Acc:** Accuracy, **MCC:** Matthews Correlation Coefficient, **AUROC:** Area Under the Receiver Operating Characteristic curve, **N5/N10/N15:** First 5/10/15 elements from N-terminal, **C5/C10/C15:** First 5/10/15 elements
from C-terminal, **N5C5/N10C10/N15C15**: First 5/10/15 elements from N-terminal as well as from C-terminal joined together.

### Model Performance on Alternate Dataset

Amino acid composition based prediction models were also developed for the alternate dataset similarly as developed for the main dataset. In this case too, the ETree based model performed best with sen of 90.23%, spc of 89.97%, acc of 90.10%, MCC of 0.80, and AUROC of 0.97 on the training dataset. On the validation dataset, the model obtained sen of 92.27%, spc of 91.75%, acc of 92.01%, MCC of 0.84, and AUROC of 0.97. The detailed result obtained using different classifiers is provided in **Table 4.** For the dipeptide composition based model too, ETree based model performed best with 90.75% sen, 88.82% spc, 89.78% acc, 0.80 MCC, and 0.96 AUROC on the training dataset. On the validation dataset, it achieved 90.72% sen, 90.21% spc, 90.46% acc, 0.81 MCC, and 0.96 AUROC. Performance obtained using various classifiers is shown in **Table 5.**

**Table 4.**
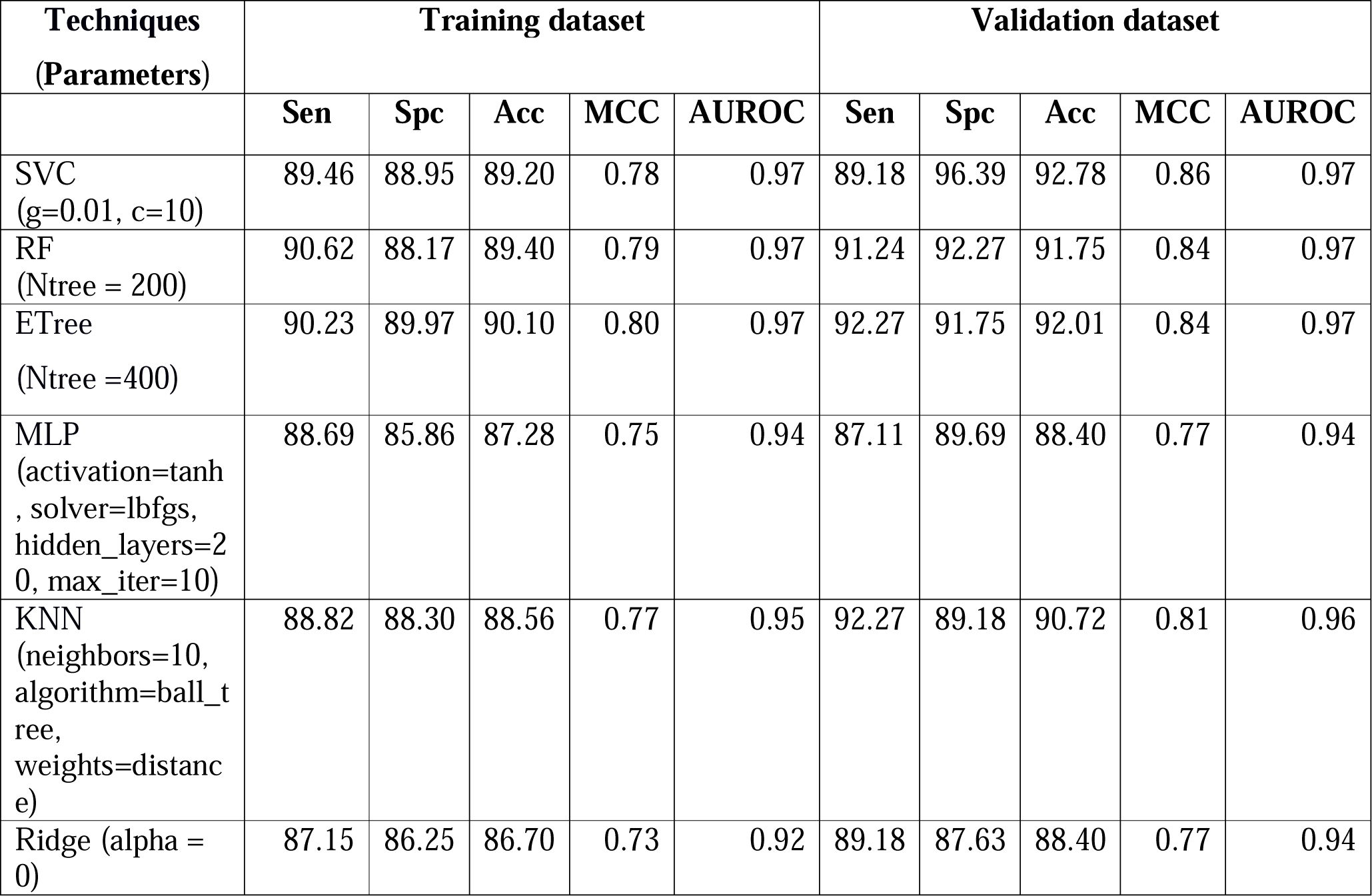
The performance of amino acid composition based models developed using different machine learning techniques on alternate dataset.

| Techniques<br>(Parameters) | Training dataset |  |  |  |  | Validation dataset |  |  |  |  |
| --- | --- | --- | --- | --- | --- | --- | --- | --- | --- | --- |
|  | Sen | Spc | Acc | MCC | AUROC | Sen | Spc | Acc | MCC | AUROC |
| SVC<br>(g=0.01, c=10) | 89.46 | 88.95 | 89.20 | 0.78 | 0.97 | 89.18 | 96.39 | 92.78 | 0.86 | 0.97 |
| RF<br>(Ntree = 200) | 90.62 | 88.17 | 89.40 | 0.79 | 0.97 | 91.24 | 92.27 | 91.75 | 0.84 | 0.97 |
| ETree<br>(Ntree =400) | 90.23 | 89.97 | 90.10 | 0.80 | 0.97 | 92.27 | 91.75 | 92.01 | 0.84 | 0.97 |
| MLP<br>(activation=tanh<br>, solver=lbfgs,<br>hidden_layers=2<br>0, max_iter=10) | 88.69 | 85.86 | 87.28 | 0.75 | 0.94 | 87.11 | 89.69 | 88.40 | 0.77 | 0.94 |
| KNN<br>(neighbors=10,<br>algorithm=ball_t<br>ree,<br>weights=distanc<br>e) | 88.82 | 88.30 | 88.56 | 0.77 | 0.95 | 92.27 | 89.18 | 90.72 | 0.81 | 0.96 |
| Ridge (alpha =<br>0) | 87.15 | 86.25 | 86.70 | 0.73 | 0.92 | 89.18 | 87.63 | 88.40 | 0.77 | 0.94 |

**Table 5.** The performance of dipeptide composition based models developed using different machine learning techniques on alternate dataset.

| Techniques<br>(Parameters) | Training dataset |  |  |  |  | Validation dataset |  |  |  |  |
| --- | --- | --- | --- | --- | --- | --- | --- | --- | --- | --- |
|  | Sen | Spc | Acc | MCC | AUROC | Sen | Spc | Acc | MCC | AUROC |
| SVC<br>(g=0.001, c=2) | 90.36 | 87.53 | 88.95 | 0.78 | 0.96 | 90.72 | 87.11 | 88.92 | 0.78 | 0.95 |
| RF<br>(Ntree = 1000) | 88.56 | 87.53 | 88.05 | 0.76 | 0.95 | 89.18 | 87.63 | 88.40 | 0.77 | 0.95 |
| ETree<br>(Ntree =400) | 90.75 | 88.82 | 89.78 | 0.80 | 0.96 | 90.72 | 90.21 | 90.46 | 0.81 | 0.96 |
| MLP<br>(activation=logistic,<br>solver=adam,<br>hidden_layers=20,<br>max_iter=10) | 87.79 | 85.22 | 86.50 | 0.73 | 0.93 | 86.08 | 88.66 | 87.37 | 0.75 | 0.94 |
| KNN<br>(neighbors=9,<br>algorithm=brute,<br>weights=distance) | 91.77 | 80.46 | 86.12 | 0.73 | 0.94 | 94.33 | 74.23 | 84.28 | 0.70 | 0.94 |
| Ridge (alpha = ) | 84.19 | 84.96 | 84.58 | 0.69 | 0.90 | 85.05 | 83.51 | 84.28 | 0.69 | 0.91 |

Similar to the main dataset, SVC based models were also developed for alternate dataset using the split amino acid composition for various patterns. The model developed using the N15C15 pattern performed best with 0.84 MCC, and 0.97 AUROC on the training dataset and 0.89 MCC, and 0.97 AUROC on the validation dataset. A detailed result for other patterns is provided in **Supplementary Table S4.** Likewise, in the case of SVC based models developed using split dipeptide composition, the model developed using the N15C15 pattern performed best. It achieved the highest MCC of 0.81, and AUROC of 0.96 on the training dataset and MCC of 0.84 and AUROC of 0.95 on the validation dataset. The result for other patterns is provided in **Supplementary Table S5.**

In the case of binary profile based models for the alternate dataset, the model developed using N15C15 pattern performed best among all models. On training dataset, sen of 88.68%, spc of 86.36%, acc of 87.43%, MCC of 0.75, and AUROC of 0.95 was obtained. On the validation dataset, the model achieved sen of 88.19%, spc of 88.16%, acc of 88.18%, MCC, of 0.76 and AUROC of 0.95. The result for all other patterns for the alternate dataset is provided in **Table 6.**

**Table 6.** The performance of SVM based model developed on alternate dataset, where models were developed using binary profile of part of peptide.

| Techniques<br>(Parameters) | Training dataset |  |  |  |  | Validation dataset |  |  |  |  |
| --- | --- | --- | --- | --- | --- | --- | --- | --- | --- | --- |
|  | Sen | Spc | Acc | MCC | AUROC | Sen | Spc | Acc | MCC | AUROC |
| N5<br>(g=0.5, c=4) | 84.50 | 83.29 | 83.39 | 0.68 | 0.92 | 88.14 | 87.63 | 87.89 | 0.76 | 0.93 |
| N10<br>(g=0.01, c=3) | 85.95 | 83.14 | 84.58 | 0.69 | 0.92 | 88.48 | 86.63 | 87.60 | 0.75 | 0.94 |
| N15<br>(g=0.01, c=3) | 85.09 | 84.74 | 84.90 | 0.70 | 0.93 | 87.50 | 85.53 | 86.49 | 0.73 | 0.94 |
| C5<br>(g=0.5, c=1) | 82.30 | 75.84 | 79.06 | 0.58 | 0.88 | 83.51 | 86.08 | 84.79 | 0.70 | 0.90 |
| C10<br>(g=0.01, c=3) | 81.04 | 79.14 | 80.11 | 0.60 | 0.88 | 80.63 | 82.56 | 81.54 | 0.63 | 0.89 |
| C15<br>(g=0.01, c=3) | 83.02 | 84.90 | 84.03 | 0.68 | 0.90 | 81.25 | 88.82 | 85.14 | 0.70 | 0.91 |
| N5C5<br>(g=0.01, c=3) | 84.88 | 81.75 | 83.31 | 0.67 | 0.91 | 86.08 | 84.54 | 85.31 | 0.71 | 0.93 |
| N10C10<br>(g=0.01, c=4) | 87.99 | 87.43 | 87.72 | 0.75 | 0.94 | 85.34 | 91.28 | 88.15 | 0.77 | 0.95 |
| N15C15<br>(g=0.01, c=1) | 88.68 | 86.36 | 87.43 | 0.75 | 0.95 | 88.19 | 88.16 | 88.18 | 0.76 | 0.95 |

#### (D) Performance on additional dataset

The performance of the prediction models developed using various input features was evaluated on an additional dataset. In the case of the main dataset, the dipeptide composition based model performed best with MCC of 0.38 and AUROC of 0.79. In the case of the alternate dataset, the amino acid composition-based model performed best with 0.70 MCC and 0.95 AUROC. The performance of models developed on other input features is provided in **Supplementary Table S6.**

### Benchmarking with existing methods

We also benchmarked the performance of our method with the existing methods on the validation dataset of both datasets (main and alternate). We observed that our model outperformed previously existing methods on both datasets, as shown in **Table 7**. We observed that other methods were predicting most of the peptides as positive i.e., indicating higher sensitivity and poor specificity in the case of the main dataset. However, our method discriminates the peptides as positive or negative with balanced sensitivity and specificity. In the case of the alternate dataset, all the methods were able to distinguish positive peptides from negative peptides with balanced sensitivity and specificity; however, the accuracy of our model among all the methods was highest. The webserver of some of the methods like ACPP, iACP were not working. Methods like iACP-GAEnSC, SAP, and Hajishraifi et al. do not provide web-based service. MLACP is predicting cell-penetrating peptide potency instead of predicting anticancer potency of a given peptide. Therefore, these methods were excluded during the benchmarking study.

**Table 7.**
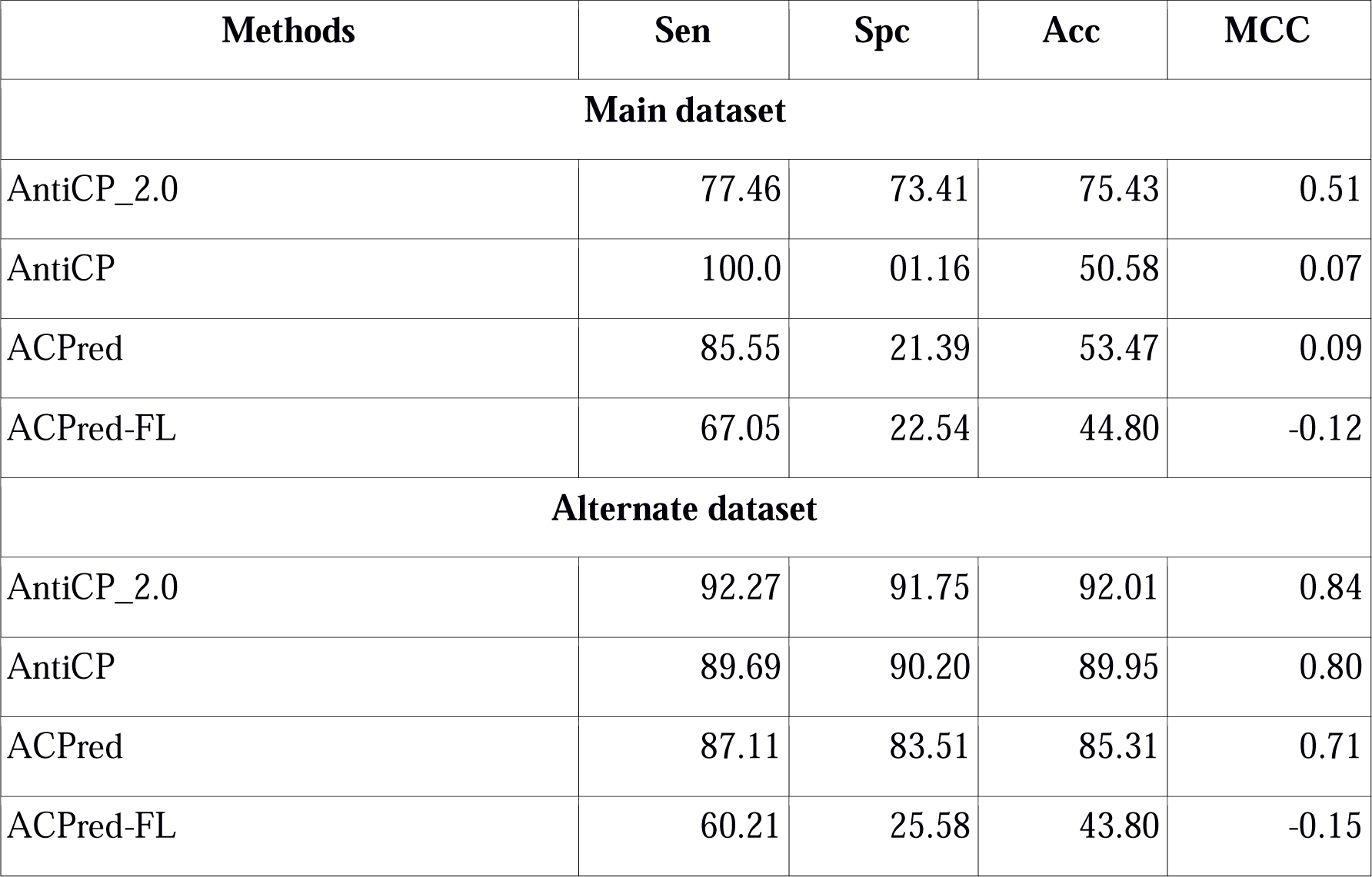
Benchmarking result of various methods on validation dataset of main and alternate dataset.

### Implementation of webserver

In the present study, for assisting experimental biologists in easy screening and designing of anticancer peptides, we have developed a web-based service “AntiCP 2.0”. We have implemented our two best models trained using two datasets i.e., main dataset and alternate dataset. “Model 1” is developed using the main dataset and will help in classifying anticancer peptides from antimicrobial peptides having activity other than anticancer. “Model 2” is developed using the alternate dataset and will help in classifying anticancer peptides from random peptides. Major modules implemented in web server includes (a) PREDICT; (b) DESIGN; (c) Protein Scan; and (iv) Download. Detail description of these modules is provided below.

a. **PREDICT:** This module predicts the anticancer potency of the submitted peptides. Users can submit multiple peptides in FASTA format in the box or can upload the file containing the same. The server will provide the result in the form of “ACP” or “non-ACP,” along with prediction score and physiochemical properties selected during submission.
b. **DESIGN:** This module allows the user to design novel ACPs with better activity. Users can enhance the activity by selecting the peptide with the best mutation and prediction score. The input is given in the form of a single line (no FASTA). The server will generate all possible mutants of the peptides with a single mutation. These mutants peptides will further get predicted by the selected model. The prediction score, along with the prediction result, will be provided based on the chosen threshold value. In the next step, the user can chose the best mutant peptides and use it further for generating new mutant peptides.
c. **Protein Scan:** In this module, the user needs to submit the protein sequence in a single line. The server will generate overlapping patterns of the protein by choosing the appropriate size. Next, based on the selected threshold value, the server will predict the anticancer potency of all the generated overlapping patterns. The result page will provide the sequence information, the prediction score, and the prediction result i.e., whether the peptide is ACP or non-ACP. This facility allows users to scan the possible anticancer region in the given protein sequence.
d. **Download:** Users can download the datasets used in this study. The sequence are present in the FASTA file format.

The webserver is freely accessible at https://webs.iiitd.edu.in/raghava/anticp2.

### Standalone

In order to serve the community, we have developed the standalone of the software in Python. Users can download the code and other required files from our GitHub account https://github.com/raghavagps/anticp2/. We have also provided a standalone facility in the form of docker technology. This standalone is integrated into our package “GPSRDocker” which can be downloaded from the site https://webs.iiitd.edu.in/gpsrdocker/ [44].

## Discussion

Peptide-based therapeutics has gained tremendous attention in the last few decades. This is reflected by the increasing number of publications in the literature, development of in silico tools and databases, etc. Due to several advantages over traditional small molecule drugs such as high specificity, less toxicity, increased half live, easy chemical modification, etc. FDA has approved several peptide-based drugs. In the same context, peptide-based anticancer drugs have shown promising effects in treating cancer [45]. Some of the peptide-based drugs for treating cancer reported in literature includes GnRH-targeting peptides [46], LY2510924 [47], ATSP-7041 [48], etc. As identification and screening of potential anticancer peptides in the wet lab are time-consuming, costly, and labour-intensive process, there is a need for advanced *in silico* tools which can do the same. Several methods have been made in the past five years for developing tools that can predict and design novel ACPs. We analysed the residue composition of ACPs and found that they are rich in residues like A, F, H, K, L, and W. It has been shown in the literature that anticancer peptides are rich in cationic residues which agrees with our study. As the order of the residues present in the peptide is strongly related to their activity, we analysed the residue preference in the ACPs. We observed that residues like F, A, and K are highly preferred at N-terminus, and L and K are preferred at C-terminus. Thus apart from the composition, the order of the residue is an important feature and might be a key feature in determining their activity. We utilized the properties of the experimentally validated ACPs present in the literature for developing various prediction models. These features include composition (amino acid and dipeptide), split composition, and binary profiles. In our study, we observed that the dipeptide composition based feature performed best among all models in the case of the main dataset. In contrast, in the case of the alternate dataset, the amino acid composition-based model performed best. In the case of the additional dataset too, dipeptide and amino acid composition based models performed best for main and alternate datasets, respectively. In the benchmarking study, we found that our model outperformed other methods.

In order to help biologists, we developed a web server and standalone app and incorporated our best models. The webserver is freely accessible and provides several facilities to the user. The server is user-friendly and compatible with multiple screens such as laptops, android mobile phones, iPhone, iPad, etc. The complete architecture of the AntiCP 2.0 is shown in **Figure 4**.

**Figure 4:**
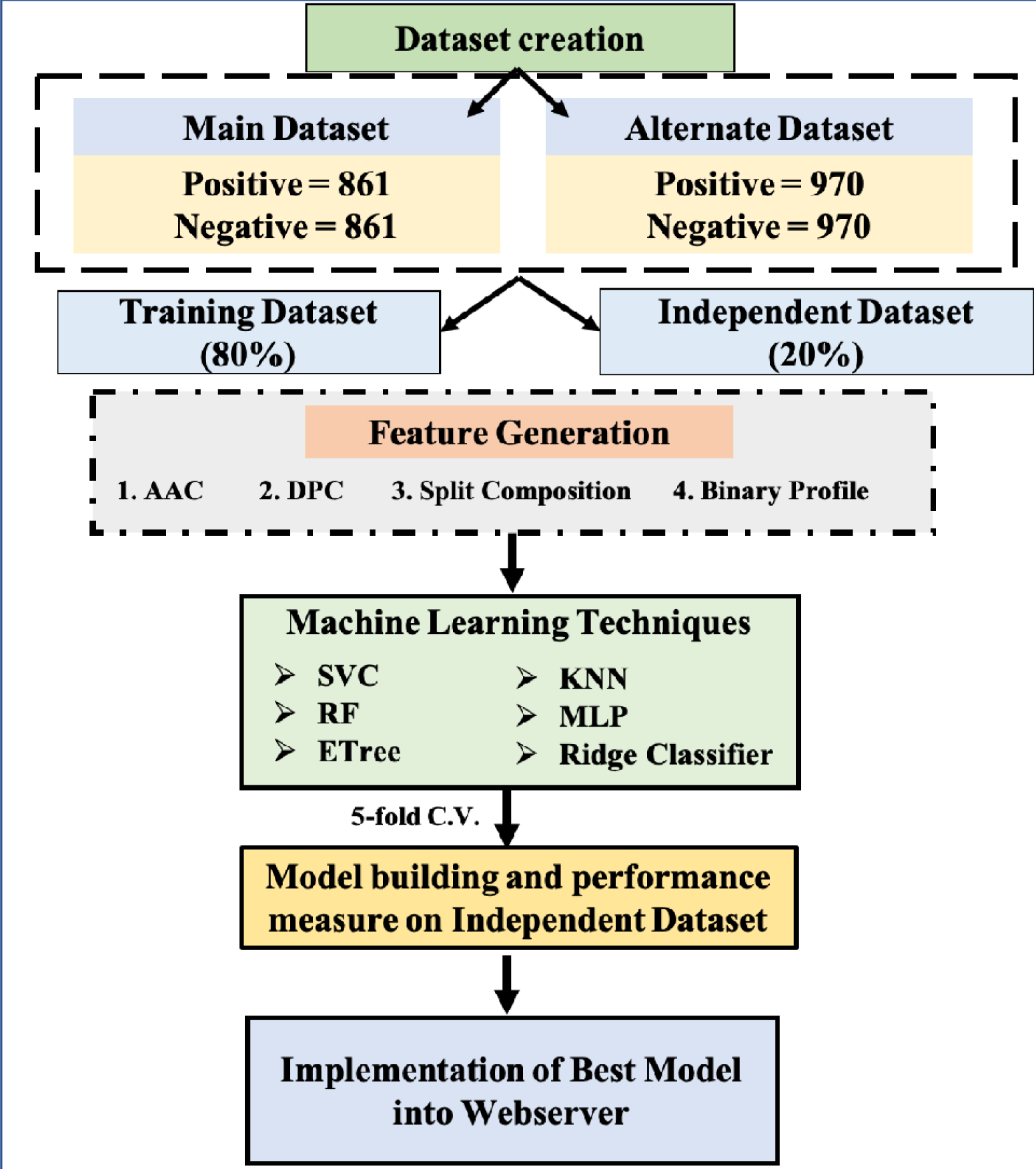
Architecture of AntiCP 2.0

## Author’s Contribution

MM, DB, and PA collected and compiled the datasets. MM, DB, and PA performed the experiments. NS, and PA developed the web interface and docker image. DB and PA developed the python based standalone software. PA, NS, and GPSR analysed the data and prepared the manuscript. GPSR conceived the idea and coordinated the project. All authors read and approved the final paper.

## Acknowledgement

Authors are thankful to J.C. Bose National Fellowship, Department of Science and Technology (DST), Government of India, and DST-INSPIRE for fellowships and the financial support.

