## Supplementary file for "AntiCP 2.0: An updated model for predicting anticancer peptides"

**^*^ Corresponding author**

Prof. G.P.S. Raghava,

Head of Department, Department of Computational Biology, Indraprastha Institute of Information Technology, Okhla Phase 3, New Delhi-110020, India.

Phone No: +91-11-26907444

**Supplementary Data**

**Table S1. Exclusive motifs of Main dataset.**

| **Sr. No.** | **Positive Motifs** | **Negative Motifs** |
| --- | --- | --- |
| 1 | KLAKK | GLW |
| 2 | LAKL | CKIK |
| 3 | LAKLA | DLV |
| 4 | AKLAK | INW |
| 5 | KKLAK | KEA |
| 6 | KLAKKL | KIKG |
| 7 | KALK | RTG |
| 8 | FAKL | AGKG |
| 9 | LAKLAK | CKIKG |
| 10 | AKLAKK | CRV |
| 11 | KLLA | GID |
| 12 | - | GLS |
| 13 | - | GLSG |
| 14 | - | TVV |

**Table S2. The performance of SVC based model developed on main dataset, where models were developed using split amino acid composition.**

| **Techniques**  (**Parameters**) | **Training dataset** | | | | | **Validation dataset** | | | | |
| --- | --- | --- | --- | --- | --- | --- | --- | --- | --- | --- |
|  | **Sen** | **Spc** | **Acc** | **MCC** | **AUROC** | **Sen** | **Spc** | **Acc** | **MCC** | **AUROC** |
| N5  (g=0.001, c=4) | 75.90 | 70.26 | 72.95 | 0.46 | 0.81 | 74.36 | 67.72 | 71.02 | 0.42 | 0.78 |
| N10  (g=0.001, c=4) | 72.43 | 68.81 | 70.61 | 0.41 | 0.78 | 76.25 | 61.35 | 68.73 | 0.38 | 0.76 |
| N15  (g=0.001, c=4) | 72.32 | 68.67 | 70.29 | 0.41 | 0.78 | 69.03 | 64.75 | 66.67 | 0.34 | 0.74 |
| C5  (g=0.001, c=4) | 69.35 | 65.37 | 67.28 | 0.35 | 0.75 | 74.55 | 65.64 | 70.12 | 0.40 | 0.78 |
| C10  (g=0.001, c=4) | 70.81 | 70.48 | 70.64 | 0.41 | 0.79 | 75.46 | 72.56 | 74.01 | 0.48 | 0.80 |
| C15  (g=0.001, c=4) | 71.59 | 69.07 | 70.19 | 0.40 | 0.76 | 73.04 | 72.34 | 72.66 | 0.45 | 0.77 |
| N5C5  (g=0.001, c=4) | 73.95 | 73.25 | 73.60 | 0.47 | 0.83 | 77.65 | 72.19 | 74.93 | 0.50 | 0.82 |
| N10C10  (g=0.001, c=4) | 71.52 | 70.75 | 71.13 | 0.42 | 0.80 | 80.00 | 69.28 | 74.62 | 0.50 | 0.84 |
| N15C15  (g=0.001, c=4) | 73.12 | 69.26 | 70.98 | 0.42 | 0.78 | 75.65 | 70.92 | 73.05 | 0.46 | 0.79 |

*** Sen:** Sensitivity, **Spc:** Specificity, **Acc:** Accuracy, **MCC:** Matthews Correlation Coefficient, **AUROC:** Area Under the Receiver Operating Characteristic curve, **N5/N10/N15:** First 5/10/15 elements from N-terminal, **C5/C10/C15:** First 5/10/15 elements from C-terminal, **N5C5/N10C10/N15C15:** First 5/10/15 elements from N-terminal as well as from C-terminal joined together.

**Table S3. The performance of SVC based model developed on main dataset, where models were developed using split dipeptide composition.**

| **Techniques**  (**Parameters**) | **Training dataset** | | | | | **Validation dataset** | | | | |
| --- | --- | --- | --- | --- | --- | --- | --- | --- | --- | --- |
|  | **Sen** | **Spc** | **Acc** | **MCC** | **AUROC** | **Sen** | **Spc** | **Acc** | **MCC** | **AUROC** |
| N5  (g=0.001, c=4) | 74.82 | 74.35 | 74.57 | 0.49 | 0.86 | 71.79 | 79.11 | 75.48 | 0.51 | 0.83 |
| N10  (g=0.001, c=4) | 74.71 | 73.95 | 74.33 | 0.49 | 0.82 | 75.00 | 69.33 | 72.14 | 0.44 | 0.81 |
| N15  (g=0.001, c=4) | 73.21 | 69.20 | 70.98 | 0.42 | 0.78 | 71.68 | 58.99 | 64.68 | 0.31 | 0.75 |
| C5  (g=0.001, c=4) | 69.35 | 67.97 | 68.63 | 0.37 | 0.79 | 70.91 | 69.94 | 70.43 | 0.41 | 0.77 |
| C10  (g=0.001, c=4) | 74.35 | 70.32 | 72.32 | 0.45 | 0.82 | 77.91 | 68.90 | 73.39 | 0.47 | 0.80 |
| C15  (g=0.001, c=4) | 71.81 | 66.26 | 68.72 | 0.38 | 0.78 | 73.91 | 62.41 | 67.58 | 0.36 | 0.76 |
| N5C5  (g=0.0001, c=2) | 73.50 | 70.58 | 72.04 | 0.44 | 0.80 | 75.88 | 73.37 | 74.63 | 0.49 | 0.83 |
| N10C10  (g=0.0001, c=2) | 71.98 | 69.22 | 70.59 | 0.41 | 0.78 | 76.97 | 68.67 | 72.81 | 0.46 | 0.81 |
| N15C15  (g=0.0001, c=2) | 70.97 | 65.80 | 68.10 | 0.37 | 0.76 | 73.91 | 63.83 | 68.36 | 0.38 | 0.77 |

*** Sen:** Sensitivity, **Spc:** Specificity, **Acc:** Accuracy, **MCC:** Matthews Correlation Coefficient, **AUROC:** Area Under the Receiver Operating Characteristic curve, **N5/N10/N15:** First 5/10/15 elements from N-terminal, **C5/C10/C15:** First 5/10/15 elements from C-terminal, **N5C5/N10C10/N15C15:** First 5/10/15 elements from N-terminal as well as from C-terminal joined together**.**

**Table S4. The performance of SVM based model developed using split amino acid composition on alternate dataset.**

| **Techniques**  (**Parameters**) | **Training dataset** | | | | | **Validation dataset** | | | | |
| --- | --- | --- | --- | --- | --- | --- | --- | --- | --- | --- |
|  | **Sen** | **Spc** | **Acc** | **MCC** | **AUROC** | **Sen** | **Spc** | **Acc** | **MCC** | **AUROC** |
| N5  (g=0.001, c=1) | 81.27 | 80.85 | 81.06 | 0.62 | 0.89 | 84.54 | 81.96 | 83.25 | 0.67 | 0.91 |
| N10  (g=0.001, c=1) | 86.08 | 84.00 | 85.07 | 0.70 | 0.92 | 86.39 | 88.95 | 87.60 | 0.75 | 0.95 |
| N15  (g=0.001, c=1) | 87.74 | 86.36 | 87.00 | 0.74 | 0.94 | 88.89 | 91.45 | 90.20 | 0.80 | 0.96 |
| C5  (g=0.001, c=1) | 79.07 | 75.19 | 77.13 | 0.54 | 0.85 | 80.41 | 77.32 | 78.87 | 0.58 | 0.88 |
| C10  (g=0.001, c=1) | 81.86 | 81.14 | 81.51 | 0.63 | 0.90 | 85.86 | 90.12 | 87.88 | 0.76 | 0.94 |
| C15  (g=0.001, c=1) | 87.55 | 85.55 | 86.47 | 0.73 | 0.94 | 91.67 | 90.79 | 91.22 | 0.82 | 0.96 |
| N5C5  (g=0.001, c=1) | 90.83 | 78.41 | 84.60 | 0.70 | 0.94 | 88.66 | 81.96 | 85.31 | 0.71 | 0.94 |
| N10C10  (g=0.001, c=1) | 91.00 | 85.57 | 88.35 | 0.77 | 0.95 | 92.15 | 90.70 | 91.46 | 0.83 | 0.97 |
| N15C15  (g=0.001, c=1) | 92.26 | 92.05 | 92.15 | 0.84 | 0.97 | 91.67 | 96.71 | 94.26 | 0.89 | 0.97 |

*** Sen:** Sensitivity, **Spc:** Specificity, **Acc:** Accuracy, **MCC:** Matthews Correlation Coefficient, **AUROC:** Area Under the Receiver Operating Characteristic curve, **N5/N10/N15:** First 5/10/15 elements from N-terminal, **C5/C10/C15:** First 5/10/15 elements from C-terminal, **N5C5/N10C10/N15C15:** First 5/10/15 elements from N-terminal as well as from C-terminal joined together.

**Table S5. The performance of SVM based model developed using split dipeptide composition on alternate dataset.**

| **Techniques**  (**Parameters**) | **Training dataset** | | | | | **Validation dataset** | | | | |
| --- | --- | --- | --- | --- | --- | --- | --- | --- | --- | --- |
|  | **Sen** | **Spc** | **Acc** | **MCC** | **AUROC** | **Sen** | **Spc** | **Acc** | **MCC** | **AUROC** |
| N5  (g=0.001, c=1) | 82.04 | 79.82 | 80.93 | 0.62 | 0.89 | 83.51 | 81.96 | 82.73 | 0.65 | 0.91 |
| N10  (g=0.001, c=1) | 86.36 | 82.29 | 84.37 | 0.69 | 0.93 | 85.86 | 86.05 | 85.95 | 0.72 | 0.93 |
| N15  (g=0.001, c=1) | 86.23 | 86.85 | 86.56 | 0.73 | 0.94 | 85.42 | 90.13 | 87.84 | 0.76 | 0.93 |
| C5  (g=0.001, c=1) | 80.62 | 72.75 | 76.68 | 0.54 | 0.87 | 83.51 | 77.32 | 80.41 | 0.61 | 0.89 |
| C10  (g=0.001, c=1) | 83.22 | 78.71 | 81.02 | 0.62 | 0.91 | 88.48 | 81.40 | 85.12 | 0.70 | 0.93 |
| C15  (g=0.001, c=1) | 86.60 | 82.95 | 84.64 | 0.69 | 0.92 | 88.19 | 80.92 | 84.46 | 0.69 | 0.93 |
| N5C5  (g=0.001, c=1) | 80.36 | 76.22 | 78.29 | 0.57 | 0.89 | 71.13 | 100.00 | 85.57 | 0.74 | 0.94 |
| N10C10  (g=0.001, c=1) | 88.27 | 86.00 | 87.16 | 0.74 | 0.95 | 89.01 | 92.44 | 90.63 | 0.81 | 0.96 |
| N15C15  (g=0.001, c=1) | 90.38 | 90.91 | 90.66 | 0.81 | 0.96 | 91.67 | 92.76 | 92.23 | 0.84 | 0.95 |

*** Sen:** Sensitivity, **Spc:** Specificity, **Acc:** Accuracy, **MCC:** Matthews Correlation Coefficient, **AUROC:** Area Under the Receiver Operating Characteristic curve, **N5/N10/N15:** First 5/10/15 elements from N-terminal, **C5/C10/C15:** First 5/10/15 elements from C-terminal, **N5C5/N10C10/N15C15:** First 5/10/15 elements from N-terminal as well as from C-terminal joined together.

**Table S6. The performance of SVM based models developed using different features on additional dataset.**

| **Features**  **(Parameters)** | **Similar Dataset** | | | | |
| --- | --- | --- | --- | --- | --- |
|  | **Sen** | **Spc** | **Acc** | **MCC** | **AUROC** |
| **Main dataset** | | | | | |
| Amino acid composition  (ETree, n_estimators=400) | 82.66 | 50.87 | 66.76 | 0.35 | 0.78 |
| Dipeptide composition  (ETree, n_estimators=100) | 80.92 | 55.49 | 68.21 | 0.38 | 0.79 |
| N10C10 Binary profile  (g=0.01, c=4) | 82.10 | 53.61 | 67.68 | 0.37 | 0.77 |
| **Alternate dataset** | | | | | |
| Amino acid composition  (ETree, n_estimators=400) | 93.30 | 75.77 | 84.54 | 0.70 | 0.95 |
| Dipeptide composition  (ETree, n_estimators=400) | 92.78 | 72.16 | 82.47 | 0.66 | 0.93 |
| N15C15 Binary profile  (g=0.001, c=1) | 89.58 | 65.45 | 76.70 | 0.56 | 0.91 |

**Sen:** Sensitivity, **Spc:** Specificity, **Acc:** Accuracy, **MCC:** Matthews Correlation Coefficient, **N15C15:** First 15 elements from N-terminal as well as from C-terminal joined together.
